# Reinforcement Learning informs optimal treatment strategies to limit antibiotic resistance

**DOI:** 10.1101/2023.01.12.523765

**Authors:** Davis T. Weaver, Eshan S. King, Jeff Maltas, Jacob G. Scott

## Abstract

Antimicrobial resistance was estimated to be associated with 4.95 million deaths worldwide in 2019. It is possible to frame the antimicrobial resistance problem as a feedback-control problem. If we could optimize this feedback-control problem and translate our findings to the clinic, we could slow, prevent or reverse the development of high-level drug resistance. Prior work on this topic has relied on systems where the exact dynamics and parameters were known *a priori*. In this study, we extend this work using a reinforcement learning (RL) approach capable of learning effective drug cycling policies in a system defined by empirically measured fitness landscapes. Crucially, we show that is possible to learn effective drug cycling policies despite the problems of noisy, limited, or delayed measurement. Given access to a panel of 15 *β* -lactam antibiotics with which to treat the simulated *E. coli* population, we demonstrate that RL agents outperform two naive treatment paradigms at minimizing the population fitness over time. We also show that RL agents approach the performance of the optimal drug cycling policy. Even when stochastic noise is introduced to the measurements of population fitness, we show that RL agents are capable of maintaining evolving populations at lower growth rates compared to controls. We further tested our approach in arbitrary fitness landscapes of up to 1024 genotypes. We show that minimization of population fitness using drug cycles is not limited by increasing genome size. Our work represents a proof-of-concept for using AI to control complex evolutionary processes.

## Introduction

Drug resistant pathogens are a wide-spread and deadly phenomenon that were responsible for nearly 5 million deaths worldwide in 2019^1^. Current projections suggest the global burden of antimicrobial resistance could climb to 10 million deaths per year by 2050^2^. In the US alone, 3 million cases of antimicrobial resistant infections are observed each year^3^. The increasing prevalence of pan-drug resistance has prompted the CDC to declare that we have entered a “post-antibiotic era”^3^. Despite this evident public health crisis, development of novel antibiotics has all but ceased due to the poor return on investment currently associated with this class of drugs^4^. Novel approaches to therapy design that explicitly take into account the adaptive nature of microbial cell populations while leveraging existing treatment options are desperately needed.

Evolutionary medicine is a rapidly growing discipline that aims to develop treatment strategies that explicitly account for the capacity of pathogens and cancer to evolve^5–11^. Such treatment strategies, termed “evolutionary therapies”, typically cycle between drugs or drug doses to take advantage of predictable patterns of disease evolution. Evolutionary therapies are often developed by applying optimization methods to a mathematical or simulation-based model of the evolving system under study^12–22^. For example, in castrate-resistant prostate cancer, researchers developed an on-off drug cycling protocol that allows drug-sensitive cancer cells to regrow following a course of treatment. Clinical trials have shown this therapy prevents the emergence of a resistant phenotype and enables superior long-term tumor control and patient survival compared to conventional strategies^23,24^.

Current methods for the development of evolutionary therapies require an enormous amount of data on the evolving system. For example, many researchers have optimized treatment by using genotype-phenotype maps to define evolutionary dynamics and model the evolving cell population^16,25–33^. For instance, Nichol *et al* modeled empirical drug fitness landscapes measured in *E. Coli* as a Markov chain to show that different sequences of antibiotics can promote or hinder resistance. In order to determine optimal drug sequences, they first constrained their system such that the population under selection evolved to a terminal evolutionary state prior to drug switching. Further, defining the Markov chain framework required exact knowledge of the high-dimensional genotype-phenotype map under many different drugs^16^. Most published methods for optimization of these models requires a complete understanding of the underlying system dynamics^15,16,34–36^. Such detailed knowledge is currently unobtainable in the clinical setting. Approaches that can approximate these optimal policies given only a fraction of the available information would fill a key unmet need in evolutionary medicine.

We hypothesize that reinforcement learning algorithms can develop effective drug cycling policies given only experimentally measurable information about the evolving pathogen. Reinforcement learning (RL) is a well-studied sub-field of machine learning that has been successfully used in applications ranging from board games and video games to manufacturing automation^34,37–39^. Broadly, RL methods train artificial intelligence agents to select actions that maximize a reward function. Importantly, RL methods are particularly suited for optimization problems where little is known about the dynamics of the underlying system. While previous theoretical work has studied evolutionary therapy with alternating antibiotics, none have addressed the problems of noisy, limited, or delayed measurement that would be expected in any real-world applications.^12–14,16,17^ Further, RL and related optimal control methods have been previously applied for the development of clinical optimization protocols in oncology and anesthesiology^21,40–45^.

In this study, we developed a novel approach to discovering evolutionary therapies using a well studied set of empirical fitness landscapes as a model system^26^. We explored “perfect information” optimization methods such as dynamic programming in addition to RL methods that can learn policies given only limited information about a system. We show that it is possible to learn effective drug cycling treatments given extremely limited information about the evolving population, even in situations where the measurements reaching the RL agent are extremely noisy and the information density is low.

## 1 Methods

As a model system, we simulated an evolving population of *Escherichia coli (E. coli)* using the well-studied fitness landscape paradigm, where each genotype is associated with a certain fitness under selection^16,26,29^. We relied on a previously described 4-allele landscape of the *E. coli β* -lacatamase gene where each possible combination of mutations had a measured impact on the sensitivity of an *E. coli* population to one of 15 *β* -lactam antibiotics^26^. We then defined 15 different fitness regimes on the same underlying genotype space, each representing the selective effect of one of 15 *β* -lactam antibiotics (**Table 1**)^26^. We used this well-studied *E. coli* model system because it is one of the few microbial cell populations for which a combinatorially complete genotype-phenotype mapping has been measured^26,29^. We extended this paradigm to procedurally generated landscapes with larger numbers of alleles as a sensitivity analysis (described in the supplemental methods). By simulating an evolving *E. coli* cell population using the described fitness landscape paradigm, we were able to define an optimization problem on which to train RL agents (**Fig 1**).

**Table 1.**
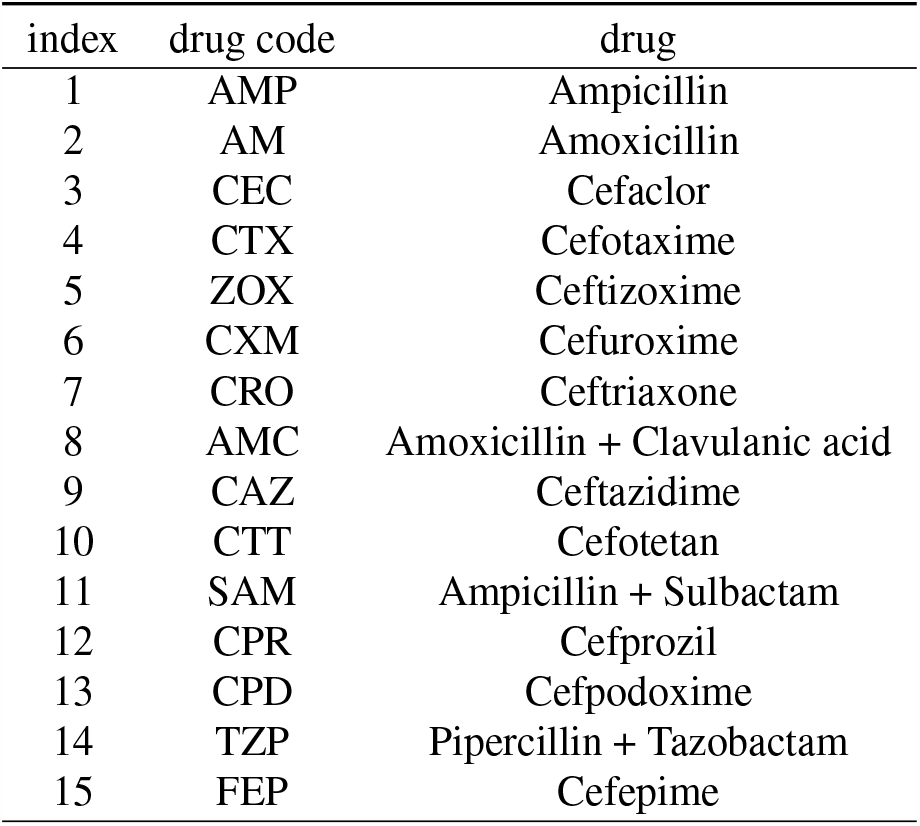
Reference codes for drugs under study.

**Figure 1.**
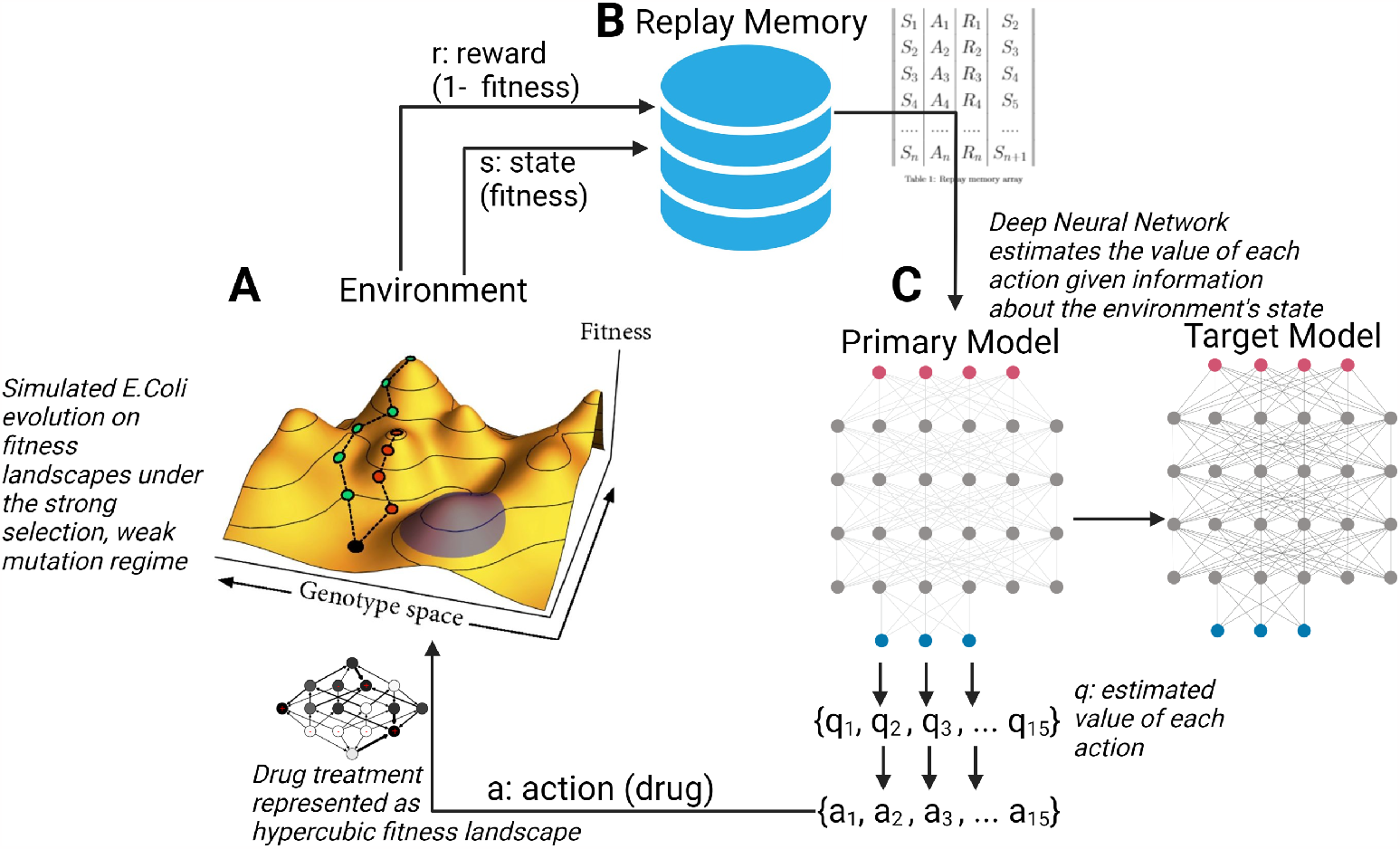
Schematic of artificial intelligence system for controlling evolving cell populations. **A:** *E. coli* population evolving on fitness landscapes under the strong selection, weak mutation evolutionary regime. At each time step, a reward signal *r* and a measure of system state *s* are sent to the replay memory structure. **B:** Replay memory array stores (s, a, r, s’) tuples where s’ is state s+1. These are then used to batch train the neural network. **C:** Deep Neural network estimates the value of each action given information about the environment’s state. The action with the largest estimated value is then applied to the evolving cell population.

### 1.1 Simulation of Evolution Using Fitness Landscapes

We use a previously described fitness-landscape based model of evolution^16,27^. In brief, we begin by modeling an evolving asexual haploid population with *N* mutational sites. Each site can have one of two alleles (0 or 1). We can therefore represent the genotype of a population using an *N*-length binary sequence, for a total of 2^*N*^ possible genotypes. We can model theoretical drug interventions by defining fitness as a function of genotype. These “drugs” can then be represented using *N*-dimensional hyper-cubic graphs (**Fig 1A**). Further, if we assume that evolution under drug treatment follows the strong selection and weak mutation (SSWM) paradigm, we can then compute the probability of mutation between adjacent genotypes and represent each landscape as a Markov chain as described by Nichol et al.^16^. With sufficiently small population size, we can then assume that the population evolves to fixation prior to transitioning to another genotype. At each time step, we sampled from the probability distribution defined by the Markov chain to simulate the evolutionary course of a single population.

### 1.2 Optimization approaches

We applied two related optimization approaches to identify effective drug cycling policies in this setting. First, we extended the Markov chain framework to formulate a complete Markov Decision Process (MDP). An MDP is a discrete-time framework for modeling optimal decision-making^34^. Critically, the system under study must be partially under the control of the decision-making agent. MDPs can be solved using dynamic programming to generate optimal policies for the defined control problem^34^. The dynamic programming algorithm requires perfect information (e.g. the complete transition matrix and instantaneous state from the MDP) in order to yield optimal policies. Next, we trained agents with imperfect information using reinforcement learning to approximate a clinical scenario where perfect information is not available. Notably, the state set, action set, and reward assignment were shared between the perfect and imperfect information conditions. The action set corresponded to the drugs available to the optimization process. We considered this system to have a finite time horizon (20 evolutionary steps in the base case). We chose a finite time horizon rather than an infinite time horizon assumption in order to more faithfully represent clinical disease courses. For our purposes, we assume that one evolutionary time step is the equivalent of one day of evolution.

#### 1.2.1 Perfect Information

The state set *S* represents all potential genotypes (16 total in our base case) that the evolving population can explore. The action set *A* corresponds to the 15 available *β* -lactam antibiotics. Finally, we define the reward set (*R*) and the set of transition probabilities (*P*) as a function of the current genotype *s* as well as the chosen action, *a* (eq 1):

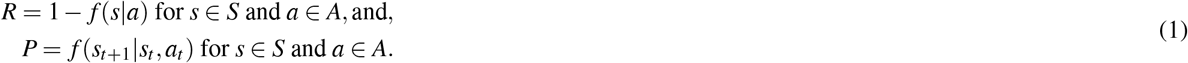

We solved the defined MDP using backwards induction, a dynamic programming approach designed to solve MDPs with finite time horizons^46^, to generate an optimal drug cycling policy for each evolutionary episode. Backwards induction is used to estimate a value function *V* (*s*) which estimates the discounted reward of being in each state *s*. Optimal policies Π(*s*) are then inferred from the value function. Throughout the remainder of the paper we will refer to this optimal drug cycling policy as the “MDP” condition.

#### 1.2.2 Imperfect Information

In order to assess the viability of developing optimal drug therapies from potentially clinically available information, we trained a Deep Q learner to interact with the evolving *E. coli* system described above. Deep Q learning is a well-studied and characterized method of reinforcement learning, and is particularly suited to situations where very little *a priori* knowledge about the environment is available^34,47^. To simulate imperfect information, we used two different training inputs to model a gradient of information loss. In the first condition, termed RL-genotype, the instantaneous genotype of the population was provided as the key training input at each time point. For this condition, the neural architecture was composed of an input layer, two 1d convolutional layers, a max pooling layer, a dense layer with 28 neurons, and an output layer with a linear activation function.

In the second condition, termed RL-fit, instantaneous population fitness of the population was provided as the key training input at each time point. The neural architecture of RL-fit was composed of a neural network with an input layer, two dense hidden layers with 64 and 28 neurons, and an output layer with a linear activation function. RL-fit takes population fitness at time *t* and one-hot encoded action at time *t*−1 as inputs and outputs Q-values. Q-values are estimates of the future value of a given action. Q-value estimates are improved by minimizing the temporal difference between Q-values computed by the current model and a target model, which has weights and biases that are only updated rarely. We used mean squared error (MSE) as the loss function.

We further explored the effect of information content on learned policy effectiveness by introducing a noise parameter. With noise active, fitness values *s* ∈ *S* that were used as training inputs were first adjusted according to:

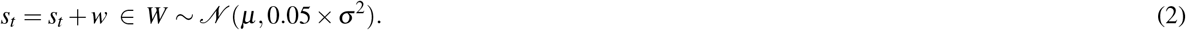

For the noise experiment,μ was set to 0 such that *σ* ^2^ = 0 would introduce no noise. We then varied *σ* ^2^ (referred to as ‘noise parameter’) from 0 (no noise) to 100 (profound loss of signal fidelity). Finally, we evaluated the performance of RL-genotype learners that were trained on delayed information to explore the viability of using outdated sequencing information to inform drug selection (described in the supplemental methods).

All code and data needed to define and implement the evolutionary simulation and reinforcement learning framework can be found at https://github.com/DavisWeaver/evo_dm. The software can be installed in your local python environment using ‘pip install git+https://github.com/DavisWeaver/evo_dm.git’. We also provide all the code needed to reproduce the figures from the paper at https://github.com/DavisWeaver/rl_cycling.

## 2 Results

In this study, we explored the viability of developing effective drug cycling policies for antibiotic treatment given less and less information about the evolving system. To this end, we developed a reinforcement learning framework to design policies that limit the growth of an evolving *E. coli* population *in silico*. We evaluated this system in a well-studied *E. coli* system for which empirical fitness landscapes for 15 antibiotics are available in the literature^26^. A given RL agent could select from any of these 15 drugs when designing a policy to minimize population fitness. We defined three experimental conditions. In the first, we solved a Markov decision process formulation of the optimization problem under study. In doing so, we generated true optimal drug cycling policies given perfect information of the underlying system (described in **Section 1.1**). In the second, RL agents were trained using the current genotype of the simulated *E. coli* population under selection (RL-genotype). Stepping further down the information gradient, RL agents were trained using only observed fitness of the *E. coli* population (RL-fit). Finally, we introduced noise into these measures of observed fitness to simulate real-world conditions where only imprecise proxy measures of the true underlying state may be available. Each experimental condition was evaluated based on its ability to minimize the fitness of the population under study in 20 time-step episodes. We compared these conditions to two negative controls; a drug cycling policy that selects drugs completely at random (which we will refer to as “random”), and all possible two-drug cycles (i.e AMP-AM-AMP-AM-AMP). We tested 100 replicates of RL-fit and RL-genotype against each of these conditions. Each replicate was trained for 500 episodes of 20 evolutionary steps (10,000 total observations of system behavior). We chose 500 episodes as the training time after extensive hyper-parameter tuning showed decreased or equal effectiveness with additional training.

### Comparison of RL drug cycling policies to negative controls

We found that both RL conditions dramatically reduced fitness relative to the random policy. In both cases, the RL conditions learned effective drug cycling policies after about 100 episodes of training and then fine-tuned them with minimal improvement through episode 500 (**Fig 2A**). As expected, RL-genotype learned a more effective drug cycling policy on average compared to RL-fit. RL-genotype had access to the instaneous state (genotype) of the evolving population, while RL-fit was only trained using a proxy measure (population fitness). We define population fitness as the instantaneous growth rate of the *E. coli* population in exponential phase. In 98/100 replicates, we observed a measurable decrease in population fitness under the learned RL-fit policy versus a random drug cycling policy (**Fig S1A**). Further, we found that the average RL-fit replicate outperformed all possible two-drug cycling policies (**Fig 2C**). RL-genotype outperformed both negative controls in all 100 replicates (**Fig 2C**). In some replicates, RL-genotype achieved similar performance compared to the MDP policy (**Fig S1D**). In addition, the distribution of performance for RL-genotype policies nearly overlapped with MDP performance (**Fig 2C**). Introduction of additional noise to the training process for RL-fit led to degraded performance. However, even with a large noise modifier, RL-fit still outperformed the random drug cycling condition. With a noise modifier of 40, RL-fit achieved an average population fitness of 1.41 compared to 1.88 for the random drug cycling condition (**Fig 2D**).

**Figure 2.**
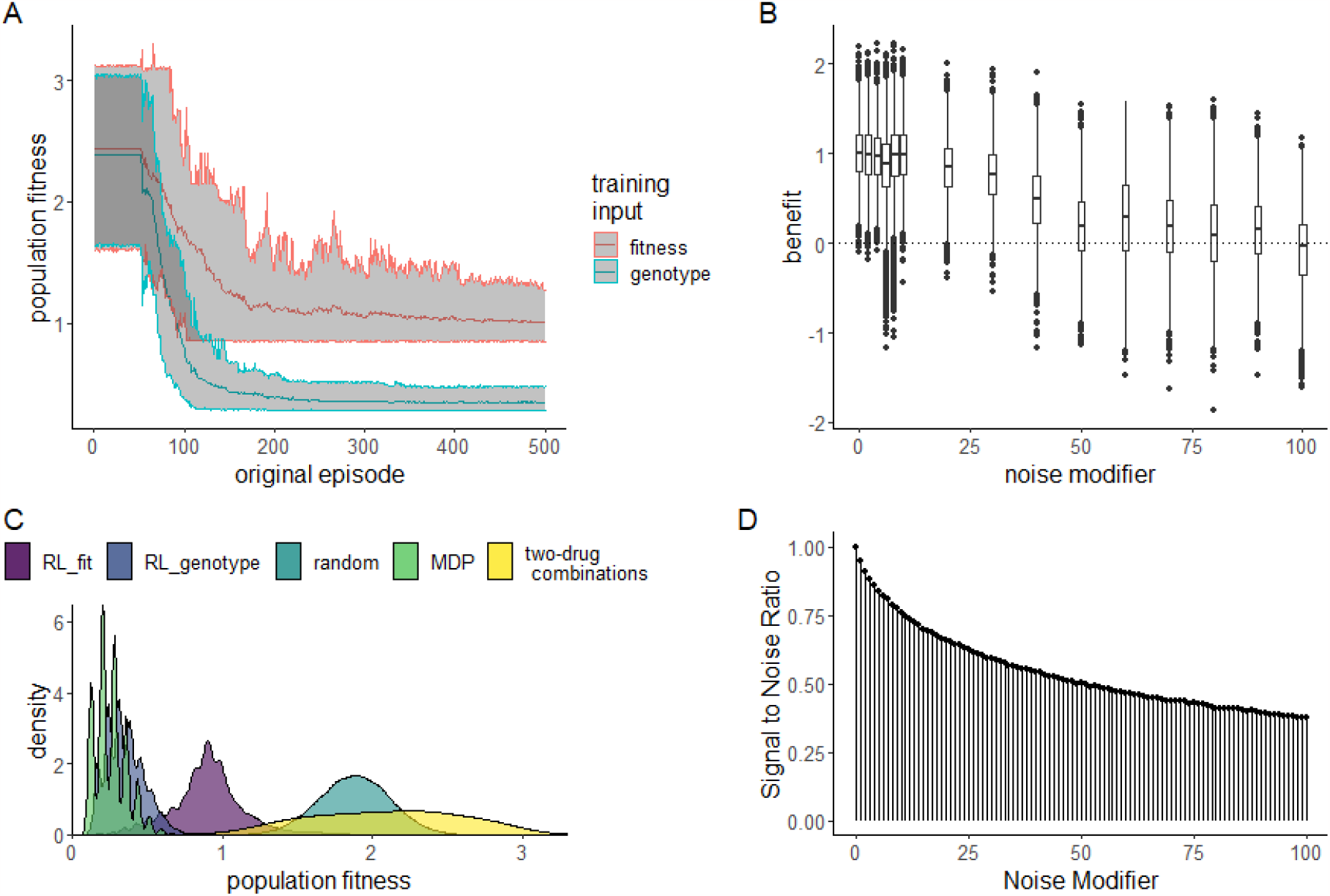
Performance of RL agents in a simulated *E. coli* system. **A:** Line plot showing the effectiveness (as measured by average population fitness) of the average learned policy as training time increases on the x-axis for RL agents trained using fitness (red) or genotype (blue). **B:** Boxplot showing the effectiveness of 10 fully trained RL-fit replicates as a function of noise. Each data point corresponds to one of 500 episodes per replicate (5000 total episodes). The width of the distribution provides information about the episode by episode variability in RL-fit performance. **C:** Density plot summarizing the performance of the two experimental conditions (measured by average population fitness) relative to the three control conditions. **D:** Signal to noise ratio associated with different noise parameters. Increasing noise parameter decreases the fidelity of the signal that reaches the reinforcement learner.

### Overview of learned drug cycling policies for RL-fit and RL-genotype

We evaluated the learned drug cycling policies of RL-fit and RL-genotype for the 15 *β* -lactam antibiotics under study. Represented drug sequences for these conditions can be found in **Table 2**. We compared these to the true optimal drug cycling policy as a reference. For this system, we show that the optimal drug cycling policy relies heavily on Cefotaxime, Ampicillin + Sulbactam, and Ampicillin (**Fig 3A**). Cefotaxime was used as treatment in more than 50% of time-steps, with Ampicillin + Sulbactam and Ampicillin used next most frequently. The optimal drug cycling policy used Cefprozil, Pipercillin + Tazobactam, and Cefaclor infrequently. The remaining drugs were not used at all. The different RL-fit replicates largely converged on a similar policy. They relied heavily on Cefotaxime and Amoxicillin + Clavulanic acid. However, they relied infrequently on Cefprozil. RL-genotype replicates also converged on a relatively conserved policy. Further, RL-genotype replicates showed a much more consistent mapping of state to action compared to RL-fit (**Fig 3B**). All optimization paradigms identified complex drug cycles that use 3 or more drugs to treat the evolving cell population. None of the tested two-drug combinations compete with policies learned by RL-genotype, and are generally out-performed by RL-fit. We show that policies that do not rely on Cefotaxime are suboptimal in this system. The three replicates that showed the least benefit compared to the random drug cycling case did not use Cefotaxime at all (**Fig 3B**). The importance of Cefotaxime is likely explained by the topography of the CTX drug landscape (**Fig S5**). More than half of the available genotypes in the CTX landscape lie in fitness valleys, providing ample opportunities to combine CTX with other drugs and “trap” the evolving population in low-fitness genotypes.

**Table 2.** Example drug sequences. Here, we show the first 10 selected drugs for representative episodes of the three top-performing replicates.

| drug sequence | replicate | condition |
| --- | --- | --- |
| CTX,AMC,CTX,CPR,CTX,CPR,CTX,CPR,CTX,CPR | 53 | RL-fit |
| CTX,CPR,CPR,CPR,CTX,CPR,CPR,CPR,CTX,SAM | 53 | RL-genotype |
| CTX,AMC,CTX,AMC,CTX,AMC,CTX,AMC,CTX,CPR | 23 | RL-fit |
| CTX,AMC,CTX,AMC,CTX,AMC,CTX,AMC,CTX,AMC | 23 | RL-genotype |
| CTX,AMC,CTX,AMC,CTX,CPR,CTX,AMC,CTX,CPR | 96 | RL-fit |
| CTX,SAM,CTX,SAM,CTX,CPR,CTX,CPR,CTX,CPR | 96 | RL-genotype |

**Figure 3.**
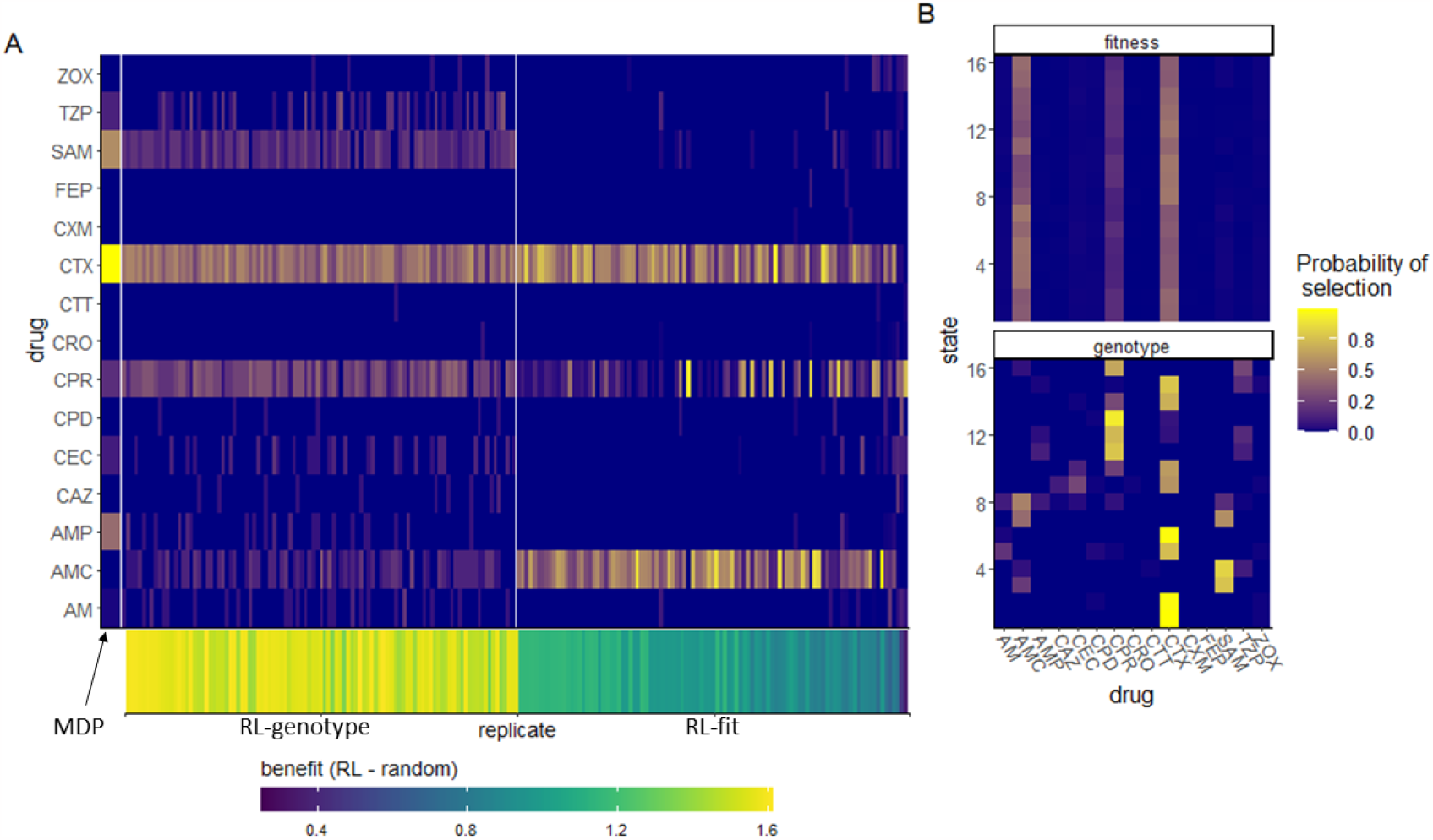
Drug cycling policies learned by RL-genotype and RL-fit. **A:** Heatmap depicting the learned policy for 100 replicates (on the x-axis) of the RL-genotype and 100 replicates of RL-fit. Far left column (enlarged) corresponds to the optimal policy derived from the MDP condition. The Y-axis describes the *β* -lactam antibiotics each RL agent could choose from while the color corresponds to the probability that the learned policy selected a given antibiotic. Bottom heatmap shows the median fitness benefit observed under the policy learned by a given replicate. **B:** Heatmap showing the average learned policy for RL-fit and RL-genotype. RL-genotype learns a more consistent mapping of state to action compared to RL-fit.

### Evolutionary trajectories observed under RL-Genotype, RL-fit, and MDP drug policies

Next, we compared the evolutionary paths taken by the simulated *E. coli* population under the MDP, RL-fit, RL-genotype, and random policy paradigms. The edge weights (corresponding to the probability of observed state transitions) of the RL-genotype and MDP landscapes show a 0.96 pearson correlation (**Fig S2**). In contrast, the edge weights of the RL-fit and MDP landscapes show a 0.82 pearson correlation (**Fig S2**). During the course of training the MDP condition, the backwards induction algorithm generated a value function *V* (*s, a*) for all *s* ∈ *S* and *a* ∈ *A*. In **Figure S2D**, we use this value function to show that certain genotypes (namely 1, 5, 6, and 13) were more advantageous to the evolving population than to the learner.These states were frequented much more often under the random drug cycling condition compared to any of the experimental conditions (**Fig S2D**). We also show that other genotypes (namely 12 and 11) were particularly advantageous for the learner compared to the evolving population. These states were frequented much more often under the experimental conditions compared to the random drug cycling condition (**Fig S2D**).

We also show that certain state transitions occur more frequently than others, independent of experimental conditions. For example, the population nearly always transitioned from genotype 5 to genotype 7 (**Fig 4**). This transition highlights the way these learned policies use drug landscapes to guide evolution. Genotype 5 (0100) is a fitness peak in most of the drug landscapes used in the learned policies, and is therefore a very disadvantageous state for the controlling agent. CTX, the most commonly used drug in all effective policies, has a slightly higher peak at genotype 7 (0110), which forces the population away from genotype 5 (**Fig S3**). As another example, the evolving population very rarely transitioned from state 1 to state 9 in the RL-fit condition. This state transition occurred commonly in the MDP and RL-genotype conditions (**Fig 4**). This difference is explained by the policies shown in **Fig 3B**. Under the RL-genotype policy, CTX was selected every time the population was in state 1 (the initial condition). The CTX landscape topography allows transition to 3 of the 4 single mutants, including state 9 (1000) (**Fig S5**). Under the RL-fit policy, CTX and AMC were used in about equal proportion when the population is in state 1. Unlike the CTX landscape, the AMC landscape topography does not permit evolution from state 1 to state 9 (**Fig S5**)).

**Figure 4.**
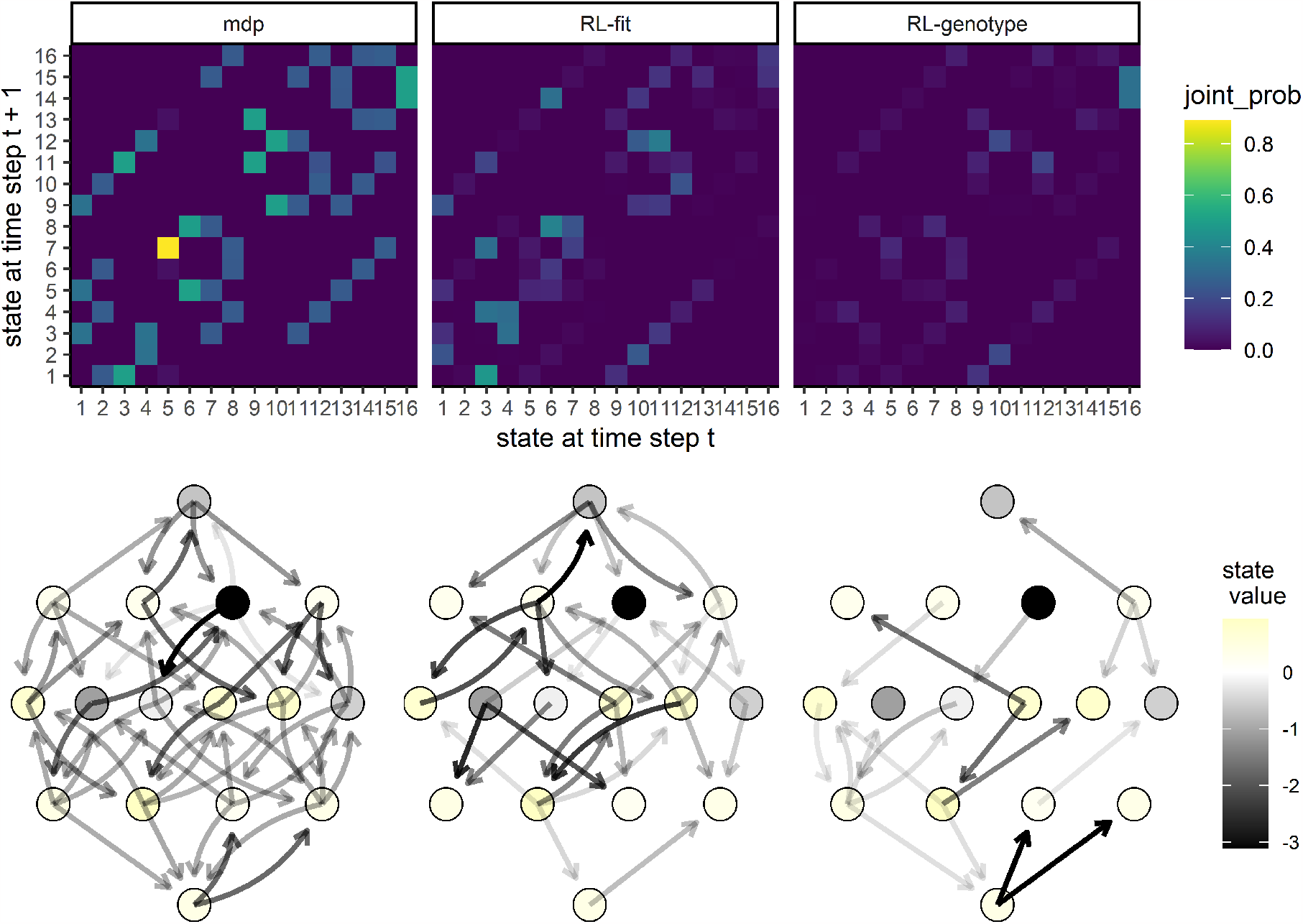
Movement of simulated *E. coli* population through the genomic landscape. **Top row:** Heatmap depicting the joint probability distribution for each state transition under the different experimental conditions. The second two show the difference in state transition probability compared to the MDP condition. **Bottom row:** Graph depicting the fitness landscape, beginning with the wild type (bottom) all the way to the quadruple mutant (top). Size of arrow depicts the frequency with which a state transition was observed under the labeled experimental condition. The color of each node corresponds with the expected value (to the learner) of being in that state. As above, the second two plots correspond to the observed difference between RL-Fit or RL-genotype and the MDP condition.

### Characteristics of selected drug policies

To better understand why certain drugs were used so frequently by RL-genotype, RL-fit, and the MDP policies, we developed the concept of an “opportunity landscape”. We computed each opportunity landscape by taking the minimum fitness value for each genotype from a given set of fitness landscapes. This simplified framework gives a sense of a potential best case scenario if the drugs in a given combination are used optimally. For example, the MDP policy relied heavily on CTX, CPR, AMP, SAM, and TZP to control the simulated *E. coli* population. The resultant opportunity landscape (**Fig 5A**) contains only a single fitness peak, with 15/16 of the genotypes in or near fitness valleys. In **Fig 5B**, we show the actual state transitions observed during evolution under the MDP policy. We also color the nodes based on the value function estimated by solving the MDP. As expected, the value function estimated by the MDP aligns closely with the topography of the opportunity landscape. There is only one genotype that the value function scores as being very poor for the learner, corresponding to the single peak in the opportunity fitness landscape (**Fig 5**). Interestingly, the opportunity landscape predicted that the population would evolve to the single fitness peak and fix. In contrast, the observed state transitions suggest that the MDP policy was able to guide the population away from that single fitness peak. A more detailed discussion of opportunity landscapes can be found in the supplemental materials.

**Figure 5.**
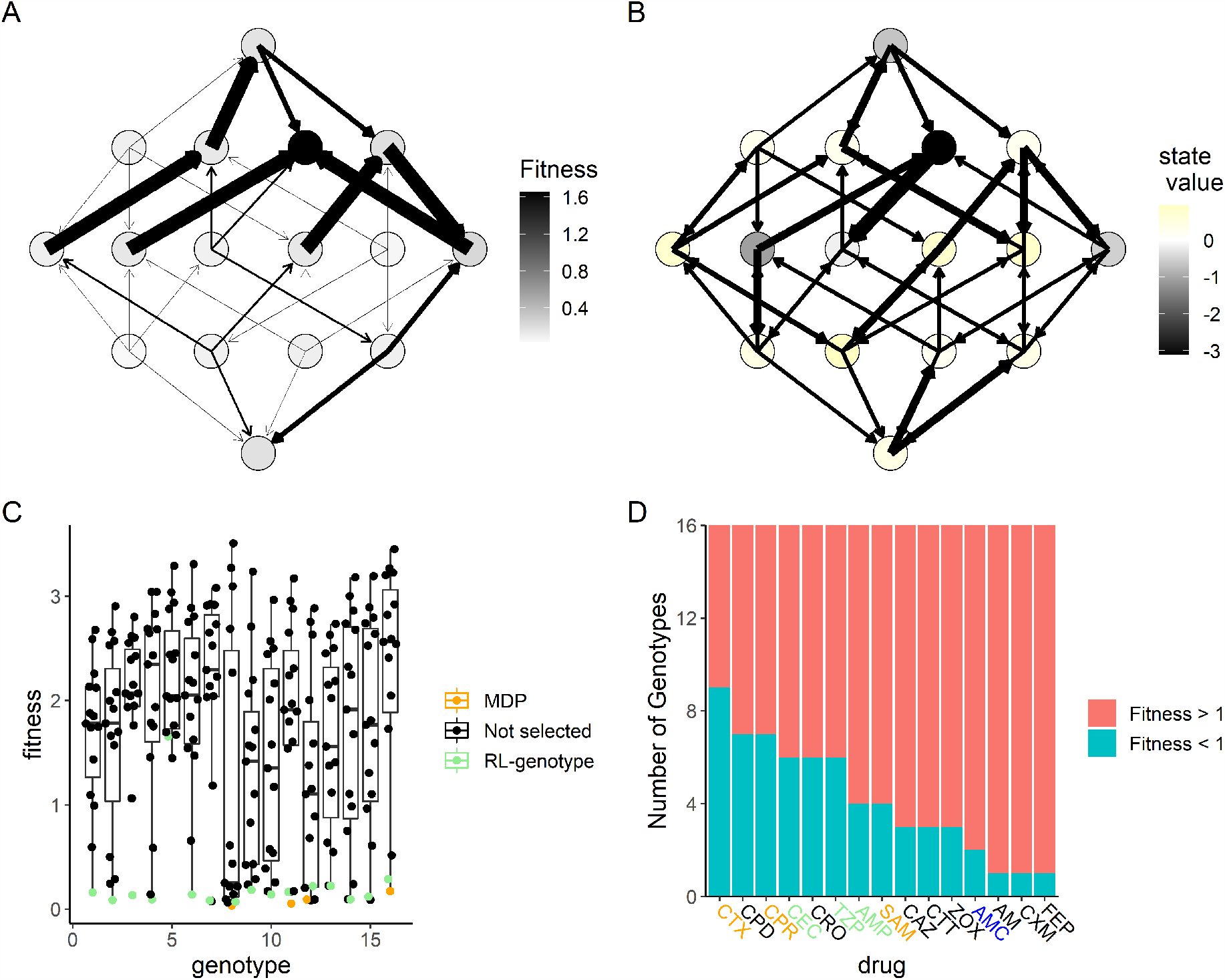
MDP value function closely matches opportunity landscape for drugs commonly used under MDP policy. Panels A and B show the 16-genotype fitness landscape under study, starting with the wild type at the top, progressing throw the single mutants, double mutants, triple mutants, and finally the quadruple mutant at the bottom. **A:** Opportunity landscape for the 5 drugs most commonly used under the MDP policy (CTX, CPR, AMP, SAM, and TZP). **B:** Observed state transitions under the MDP policy. The node color corresponds to the value function estimated by solving the MDP. Lower values correspond to states the MDP policy attempts to avoid while higher values correspond to states the MDP policy attempts to steer the population. **C:** Scatter Plot showing the distribution of fitness with respect to genotype for the 15 *β* -lactam antibiotics under study. The drug selected by RL-genotype in a given genotype is highlighted in light blue. In cases where the MDP selected a different drug than RL-genotype, that drug is highlighted in orange. **D:** Number of genotypes with fitness above or below 1 for each drug under study. Drugs that are used by both the MDP and RL-genotype are highlighted in orange. Drugs that are used by only the MDP are highlighted in green. Drugs that are used by only RL-genotype are highlighted in blue.

We also show that both the MDP and RL-genotype conditions select the drug with the lowest fitness for most genotypes (**Fig 5C**). There are a few notable exceptions to this rule, which highlight RL-genotype’s capacity for rational treatment planning. A greedy policy that selects the lowest drug-fitness combination for every genotype would select Amoxicillin (AM) when the population is identified as being in genotype 5. The AM drug landscape then strongly favors transition back to the wild-type genotype (state 1). From state 1, most available drugs encourage evolution back to the genotype 5 fitness peak. As we see in **Fig 5B**, state 5 is by far the least advantageous for the learner. The greedy policy therefore creates an extremely disadvantageous cycle of evolution. In fact, none of the tested policies rely heavily on AM in state 5 (**Fig 3B**), instead taking a fitness penalty to select Cefotaxime (CTX). The CTX drug landscape encourages evolution to the double mutant, which has access to the highest value areas of the landscape. Finally, we rank drug landscapes based on the number of genotypes with a fitness value < 1 (**Fig 5D**). Based on the defined reward function, these genotypes would be considered advantageous to the learner. We show that drugs identified as useful by the optimal policy or RL-genotype tend to have more advantageous genotypes in their drug landscape. The only two highly permissive landscapes (CPD, CPR) that aren’t used have extremely similar topography to CTX, which most policies were built around.

### Impact of landscape size and measurement delay

To understand the impact of larger fitness landscapes on the ability of our method to develop effective policies, we simulated random correlated landscapes of size *N* alleles, from *N* = 4 to *N* = 10, representing a range of 16 to 1024 genotypes. Using a previously described technique, we tuned the correlation between landscapes to generate a range of collateral resistance and collateral sensitivity profiles^27^. We found that reinforcement learners trained on fitness and genotype were able to outperform the random cycling control across a wide range of landscapes sizes (**Fig. 6A**). RL-genotype policies consistently outperformed RL-fit policies, which outperformed random drug cycling policies. Further, the MDP-derived optimal policy achieved similar performance on larger landscapes compared to smaller ones, suggesting that increasing genome size does not make drug cycling for evolutionary control less feasible. As we saw in the empirical landscapes condition, the RL-genotype and MDP policies make use of many available drugs to steer the population, while the RL-fit condition tends to identify a different policy optima that relies on 2 or 3 drugs (**Fig 6D**). Finally, we investigated the state values for the largest landscape (*N* = 10), finding that the state value space, and therefore policy space, is rugged, with many peaks and valleys (**Fig. 6C**).

**Figure 6.**
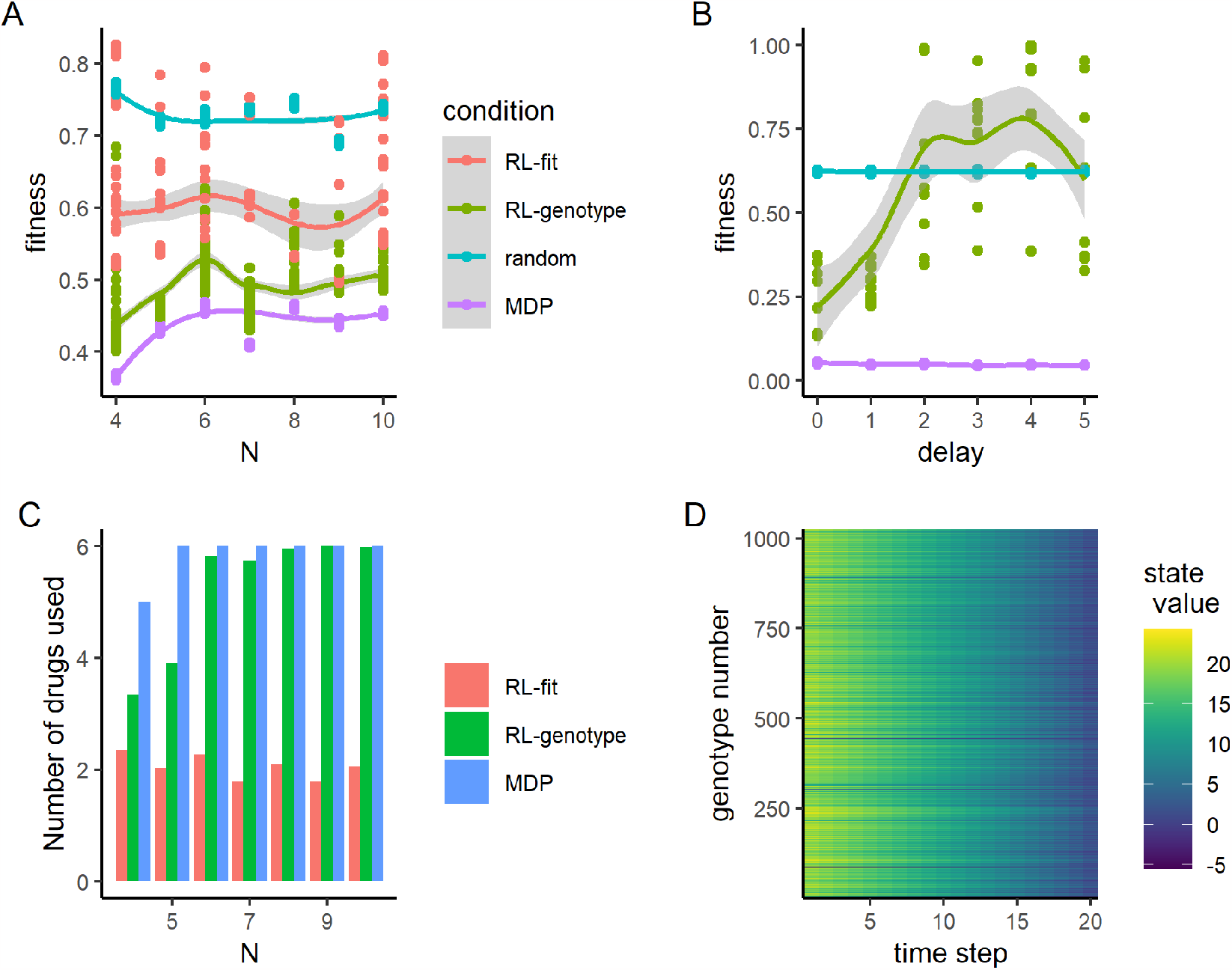
Reinforcement learners learn improved policies, independent of landscape size. **A:** Line plot describing the relationship between the number of alleles modeled and the average fitness observed under different policy regimes. The total number of states in a landscape is given by 2^*N*^. Each point represents the average fitness of a population under control of an agent trained on the set of landscapes for 500 episodes. The same set of landscapes was used for each condition. **B:** Line plot describing the relationship between measurement delay and the observed fitness under different policy regimes. Delay only affects the RL_genotype policy (not the MDP or random conditions). RL_genotype still learned effective policies with a measurement delay of one time step. Performance decayed substantially with additional measurement delays. **C:** Average number of drugs used by final policy under three experimental conditions (RL-fit, RL-genotype, MDP) as a function of *N*. **D:** Heatmap showing the value function learned by solving the MDP for the *N* = 10 landscapes. Y-axis corresponds to numerically encoded genotype and X-axis corresponds to the time step within a given episode. Bright cells correspond to genotypes that were advantageous to the agent while dark cells correspond to genotypes that were disadvantageous to the agent. The value space is rugged, with many peaks and valleys.

Time delays between when information is sampled from a system and when action is taken based on that information may be unavoidable in real-world applications. To understand how the practical limitation of time delays impacts our approach, we next tested the effect of measurement delays on the ability of reinforcement learners to generate effective policies. In this study, the delay parameter *d* controlled the “age” of the information available to the learner. With a delay of 0 (*d* = 0) time steps, the learner had completely up to date information about system state. If *d* = 5, the learner was using genotype information from time *t* to inform an action taken on time *t* + 5. We tested a set of delay parameters *d* ∈ 0, 5. For this experiment, we tested only the RL-genotype condition against the random and MDP conditions, arguing that growth rate estimates are much easier to obtain compared to sequencing, and thus measurement delays are less likely to be a practical limitation for the RL-fit condition. We did not include a delay for the MDP condition, preferring to leave it as a perfect information comparison group. We found that when *d <*= 1, average performance of the RL-genotype condition was equivalent to that observed in the base case. If *d >* 1, the performance of RL-genotype decreased to worse than or similar to the random condition (**Fig. 6B**).

## 3 Discussion

The evolution of widespread microbial drug resistance is driving a growing public health crisis around the world. In this study, we show a proof of concept for how existing drugs could be leveraged to control microbial populations without increasing drug resistance. To that end, we tested optimization approaches given decreasing amounts of information about an evolving system of *E. coli*, and showed that it is possible to learn highly effective drug cycling policies given only empirically measurable information. To accomplish this, we developed a novel reinforcement learning approach to control an evolving population of *E. coli in silico*. We focused on 15 empirically measured fitness landscapes pertaining to different clinically available *β* -lactam antibiotics (**Table 1**). In this setting, RL agents selected treatments that, on average, controlled population fitness much more effectively than either of the two negative controls. We showed that RL agents with access to the instantaneous genotype of the population over time approach the MDP-derived optimal policy for these landscapes. Critically, RL agents were capable of developing effective drug cycling protocols even when the measures of fitness used for training were first adjusted by a noise parameter. This suggests that even imperfect measurements of an imperfect measure of population state (the kind of measurements we are able to make in clinical settings) may be sufficient to develop effective control policies. We also show that RL or MDP-derived policies consistently outperform simple alternating drug cycling policies. Next, we performed a sensitivity analysis to show how RL agents can outperform random controls even with varying landscape size and measurement delay up to one day. Finally, we introduced the concept of the “Opportunity Landscape”, which can provide powerful intuition into the viability of various drug combinations.

Our work expands a rich literature on the subject of evolutionary control through formal optimization approaches. Our group and others have developed and optimized perfect information systems to generate effective drug cycling policies^12,13,15,17,18^. Further, a limited number of studies have used RL-based methods for the development of clinical optimization protocols^21,40–43,45^. These studies have been limited so far to simulated systems, including a recent study that introduced Celludose, a RL framework capable of controlling evolving bacterial populations in a stochastic simulated system^44^.

Much like the studies noted above, we show that AI or MDP-based policies for drug selection or drug dosing dramatically outperform sensible controls in the treatment of an evolving cell population. We extend this literature in three key ways. To our knowledge, ours is the first optimization protocol capable of learning effective drug cycling policies using only observed population fitness (a clinically tractable measure) as the key training input. Importantly, the reinforcement learners have no prior knowledge of the underlying model of evolution. Second, we grounded our work with empirically measured fitness landscapes from a broad set of clinically relevant drugs, which will facilitate more natural extension to the bench. Third, we tested our approach in fitness landscapes of up to 1024 genotypes, by far the largest state space that has been evaluated in the treatment optimization literature. We show that minimization of population fitness using drug cycles is not limited by increasing genome size.

There are several limitations to this work which bear mention. We assume that selection under drug therapy represents a strong-selection and weak mutation regime in order to compute transition matrices for our models. While this is likely true in most cases, it is possible that other selection regimes emerge in cases of real world pharmacokinetics or spatial regimes where the drug concentration fluctuates dramatically^48,49^. In addition, we chose to keep drug concentration constant throughout are analysis, largely owing to the lack of robust empirical data linking genotype to phenotype under dose varying conditions (sometimes called a fitness seascape)^50^. As more empirical fitness seascape data becomes available, a natural extension would be to explore the efficacy of the RL system in controlling a population by varying both drug and dose.

While we present the most extensive genotype-phenotype modeling work to date on this subject, we still only modeled the effect of mutations at up to 10 genotypic positions. The real *E. coli* genome is approximately 5×10^6^ base pairs^51^. The evolutionary landscape for living organisms is staggeringly large, and not tractable to model *in silico*. It is possible that empirical measures of fitness like growth rate or cell count may not provide a robust enough signal of the underlying evolutionary state on real genomes. *In vitro* implementations of reinforcement learning-based drug cycle optimization systems are needed to address this potential shortcoming. Another potential alternative would be to use the comparatively low-dimensional phenotype landscape of drug resistance^52^.

In this work, we present a novel reinforcement-learning framework capable of controlling an evolving population of *E. coli in silico*. We show that RL agents stably learn multi-drug combinations that were state specific and reliably out-performed a random drug cycling policy as well as all possible two-drug cycling policies. We also highlight key features of the types of drug landscapes that are useful for the design of evolutionary control policies. Our work represents an important proof-of-concept for AI-based evolutionary control, an emerging field with the potential to revolutionize clinical medicine.

## Acknowledgements

This work was made possible by the National Institute of Health (5R37CA244613-03, 5T32GM007250-46, and T32CA094186) and American Cancer Society (RSG-20-096-01). **Figure 1** was created with BioRender.com.

## Supplemental Materials

In the following please find the supplemental materials for the manuscript entitled: “Reinforcement Learning informs optimal treatment strategies to limit antibiotic resistance”

### 1.1 Procedurally generated drug landscapes

The 4 curated amino acid substitutions used to generate fitness landscapes in this study likely represent the most important mutations in the evolution of drug resistance to *β* -lactam antibiotics in this model system. However, there may be other “off-landscape” mutations in genes such as drug efflux pumps that also significantly impact resistance. Furthermore, other organisms or drug combinations may demand much larger landscapes to effectively predict evolution^53,54^. To evaluate the feasibility of reinforcement learning based drug cycle optimization in larger state spaces, we generated correlated landscapes with arbitrary *N* (where *N* is the number of alleles), following a procedure adapted from previous work^27^. We first generated an index landscape *L* by sampling a vector of length 2^*N*^ (where *N* is the number of alleles) from a uniform distribution with range (−1, 1). We then introduced epistasis by applying a gaussian noise vector (with *μ* = 0 and *σ* = 0.5) to each element of the vector. This process is traditionally referred to as a “rough Mt Fuji.” Next, we generated a set of *n* correlated landscapes *L*_*c*_ with correlations to the original landscape ranging from −1 (perfectly anti-correlated) to 1 (perfectly correlated). Briefly, we generate a Gaussian random vector (with zero mean and variance) of length 2^*N*^. We subtract from this vector its projection onto the original landscape vector *L*, making our new vector orthogonal to *L*. It then follows that any vector *L*_*c*_ is a linear combination of *L* and our new orthogonal vector. In practice, anti-correlated drug landscapes display striking collateral sensitivity, while correlated drug landscapes display collateral resistance. From the set of correlated landscapes *L*_*c*_, we selected a subset of landscapes demonstrating the full range of correlations to ensure that collateral sensitivity and collateral resistance were both present.

### 1.2 Measurement Delay

In this sensitivity analysis, we aimed to assess the viability of RL-based drug cycling policies in a setting where actions are taken based on “out-of-date” information. For example, if DNA sequencing takes multiple days to process, drugs applied based on that data would be reacting to a version of the population that no longer exists. We therefore defined a delay parameter *d* which controlled the number of time steps removed the action was from the measurement of environmental state. Put another way, *s*_*t*−*d*_ informed *a*_*t*_. If *d* = 0 (as in our base case), *s*_*t*_ informed *a*_*t*_.

We limited this analysis to the RL-genotype condition. Our rationale was that fitness or growth rate measurements are much easier to obtain and such delays wouldn’t be as common, even in complex *in vitro* settings. We hypothesized that out-of-date state vectors would lack sufficient information content to effectively inform the reinforcement learning agents.

### 1.3 Hyperparameter tuning

We varied key hyperparameters one at a time in order to identify optimal values to promote learning in this setting. Parameter ranges and the selected value are shown in **Table S1**. Due to the long run-times of the training process, we were unable to make use of more formal hyper-parameter optimization approaches. Future work will increase the efficiency of training reinforcement learners in this setting, opening up a number of interesting follow-on studies.

**Table S1.**
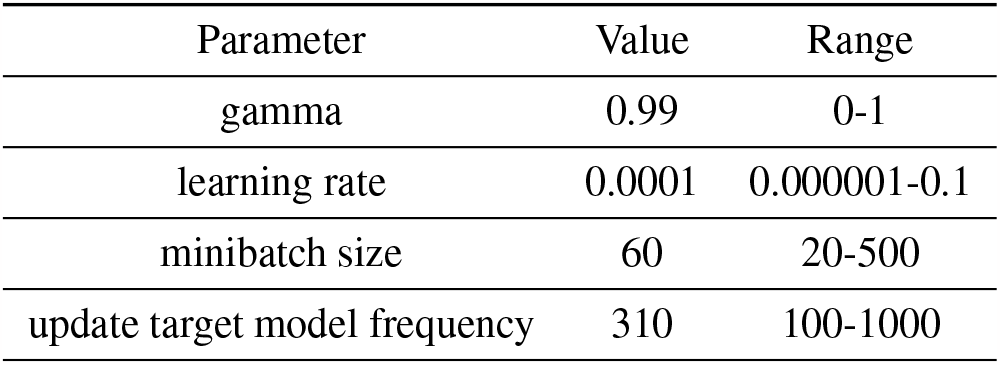
Key Hyperparameters for reinforcement learner

| Parameter | Value | Range |
| --- | --- | --- |
| gamma | 0.99 | 0-1 |
| learning rate | 0.0001 | 0.000001-0.1 |
| minibatch size | 60 | 20-500 |
| update target model frequency | 310 | 100-1000 |

### 1.4 Additional performance data for RL agents

As mentioned in the main text, we tested both the RL-fit and RL-genotype conditions 100 times each. In **Fig S1**, we show the perfomance of all 100 RL-fit and RL-genotype replicates. In 98/100 replicates, RL-fit outperformed the random drug cycling case (**Fig S1A**). The very best RL-fit replicates still fell short of the MDP-derived optimal policy (**Fig S1B**). In all 100 replicates, RL-genotype outperformed the random drug cycling case (**Fig S1C**). RL-genotype performance approached the performance of the optimal policy (**Fig S1D**).

**Figure S1.**
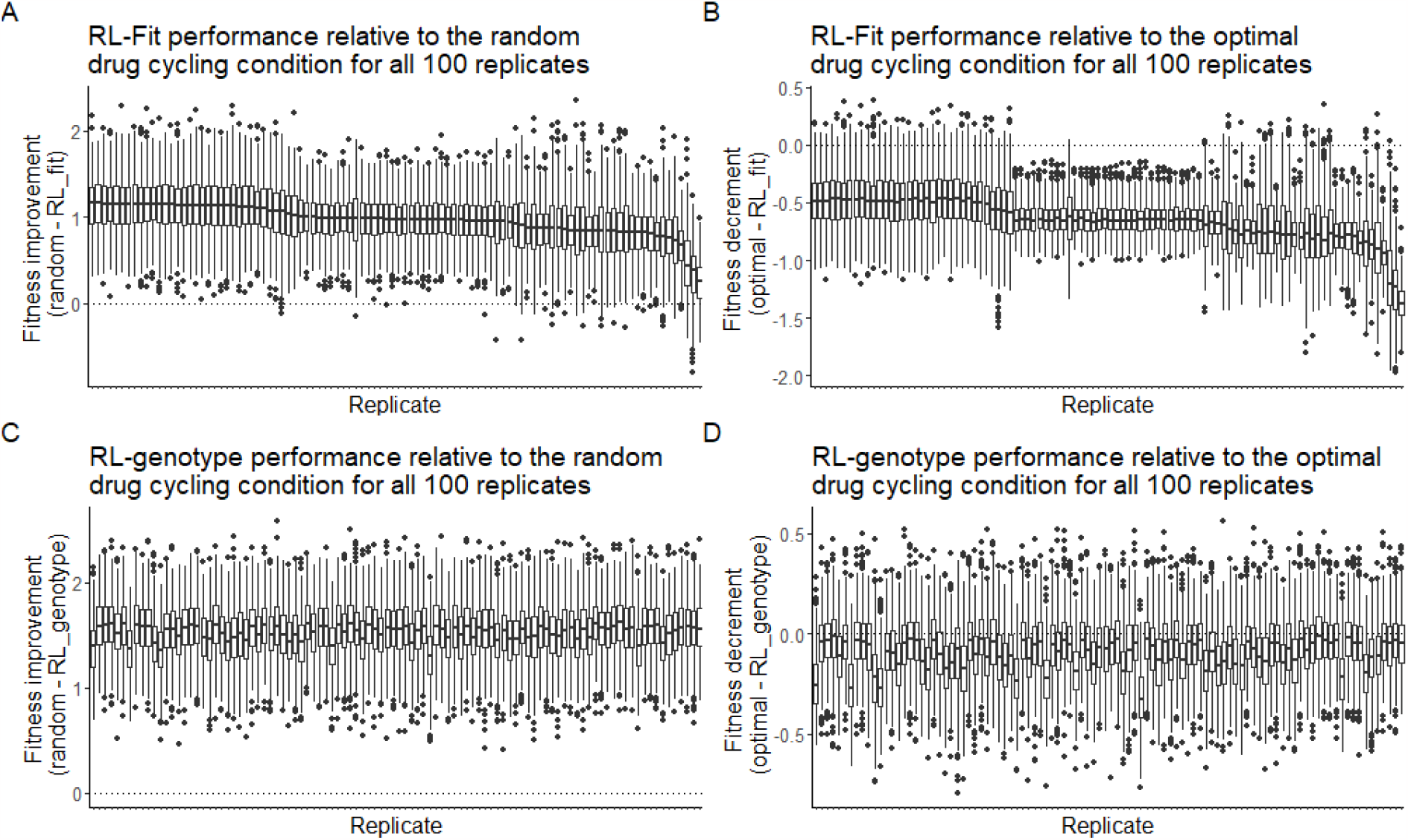
Performance of RL-fit and RL-genotype for each replicate. **A:** Fitness observed under RL-fit policy compared to random drug cycling condition. **B:** Fitness observed under RL-fit policy compared to fitness observed under optimal policy. **C:** Fitness observed under RL-genotype policy compared to random drug cycling condition. **D:** Fitness observed under RL-genotype policy compared to fitness observed under optimal policy.

### 1.5 Additional evolutionary trajectory data

We compared the state transition frequencies observed under different policy regimes, where a state transition is defined as the population evolving from genotype *s*_*a*_ to genotype *s*_*b*_. We also show the frequency with which each state was visited under different conditions. Policy performance is closely tied to the frequency with which state 5 (a high fitness genotpye in nearly all drugs) is visited. (**Fig S2**).

**Figure S2.**
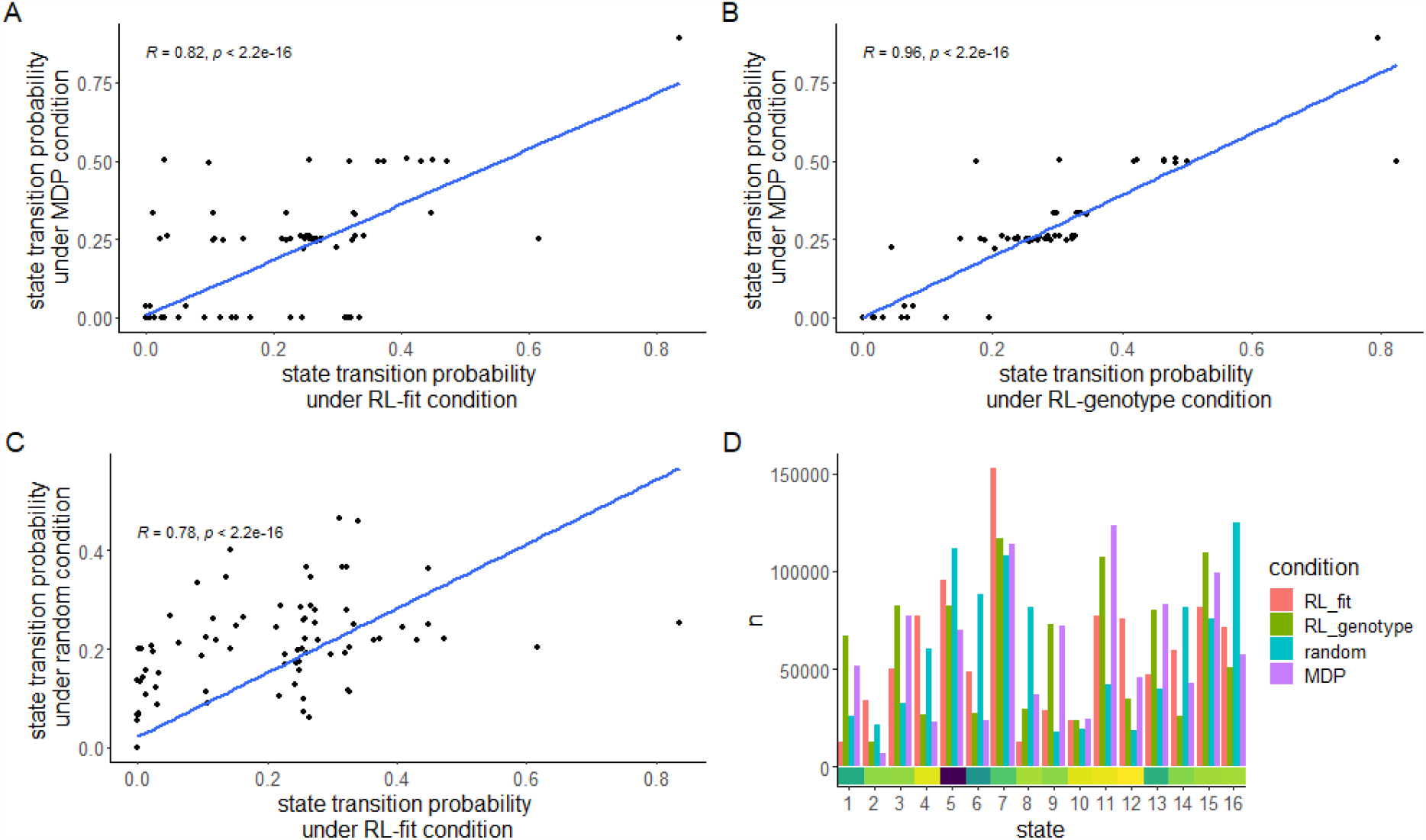
Comparison of evolutionary trajectories seen under different regimes. **A-C:** Selected Pairwise comparisons of state transition frequency under different experimental conditions. State transition frequency is nearly identical for the RL-genotype and MDP conditions (R=0.96). In contrast,state-transition-frequency for the RL-fit and MDP conditions are related but less strongly correlated (R=0.82). As expected, state transition frequency were least similar between the RL-fit and random conditions (R=0.78). **D:** Bar chart comparing the frequency that states are observed under different experimental conditions. The value of each state (to the learner) is highlighted for each state by the bottom heatmap. High value states are observed more frequently in RL-fit, RL-genotype, and MDP conditions compared to the random condition.

### 1.6 Opportunity Landscapes

We define an opportunity landscape to be the most optimistic combination of *n* landscapes, formed by taking the minimum possible fitness at each genotypic position. This construct can help us better understand how the learner uses different combinations of drugs to maintain the evolving population at extremely low fitness values.

**Figure S3.**
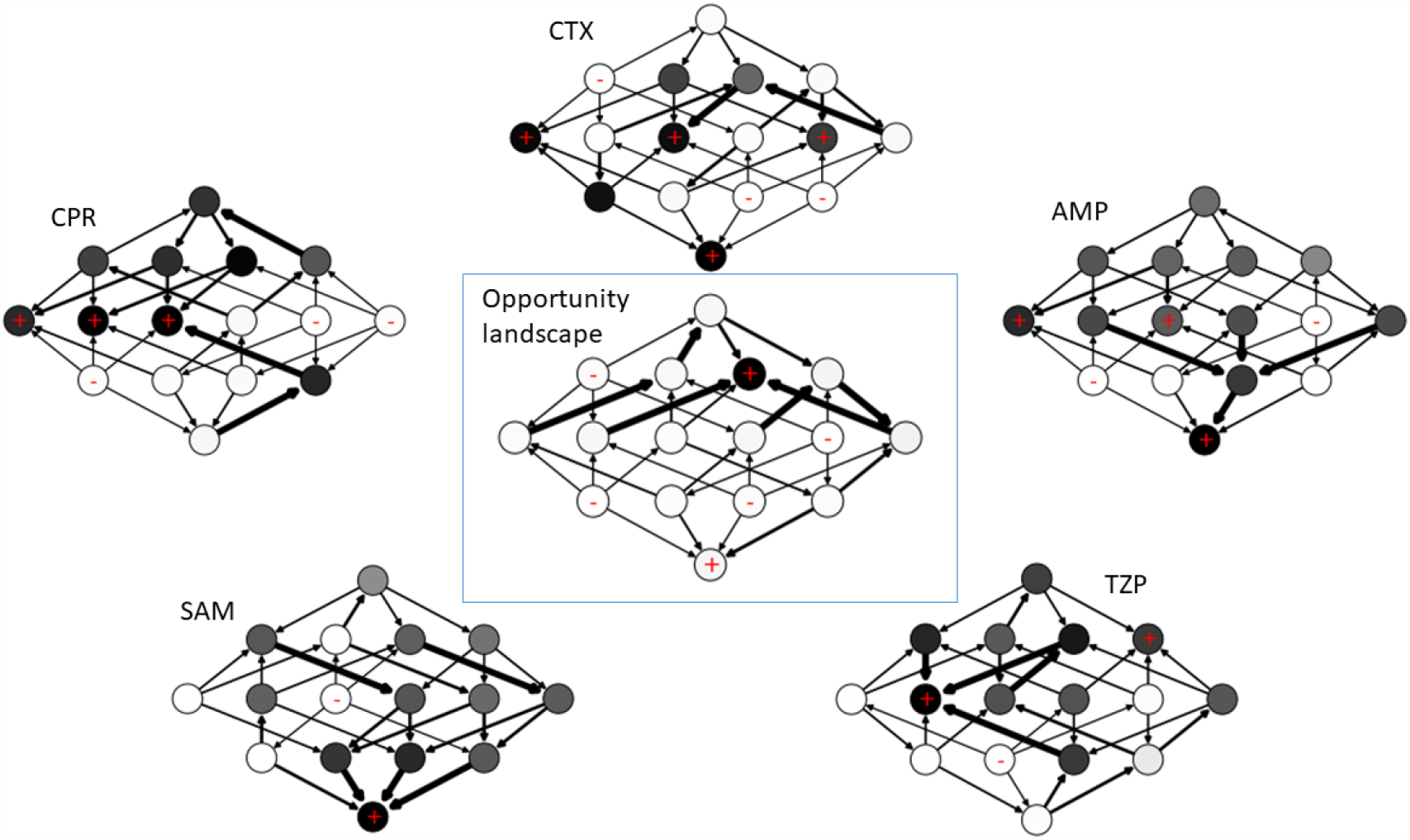
Opportunity Landscape for MDP-derived policy. Opportunity landscape is an optimistic combination of 5 empirically measured drug landscapes. Just 1/16 genotypes is near a fitness peak on the opportunity landscape, helping to explain the extremely low fitness observed in the simulated E. coli population when the MDP-derived policy is applied.

**Fig. S3** describes the opportunity landscape discovered by the MDP condition. As noted in the main text, the MDP primarily uses 5 drugs (CTX, CPR, AMP, SAM, and TZP) in combination to trap the evolving population of E. coli at extremely low fitness genotypes. In the combined opportunity landscape, just one genotype (0100) had a high fitness in all 5 drugs. As expected, the opportunity landscape closely matches the value function estimated by the MDP (**Fig 4**).

**Figure S4.**
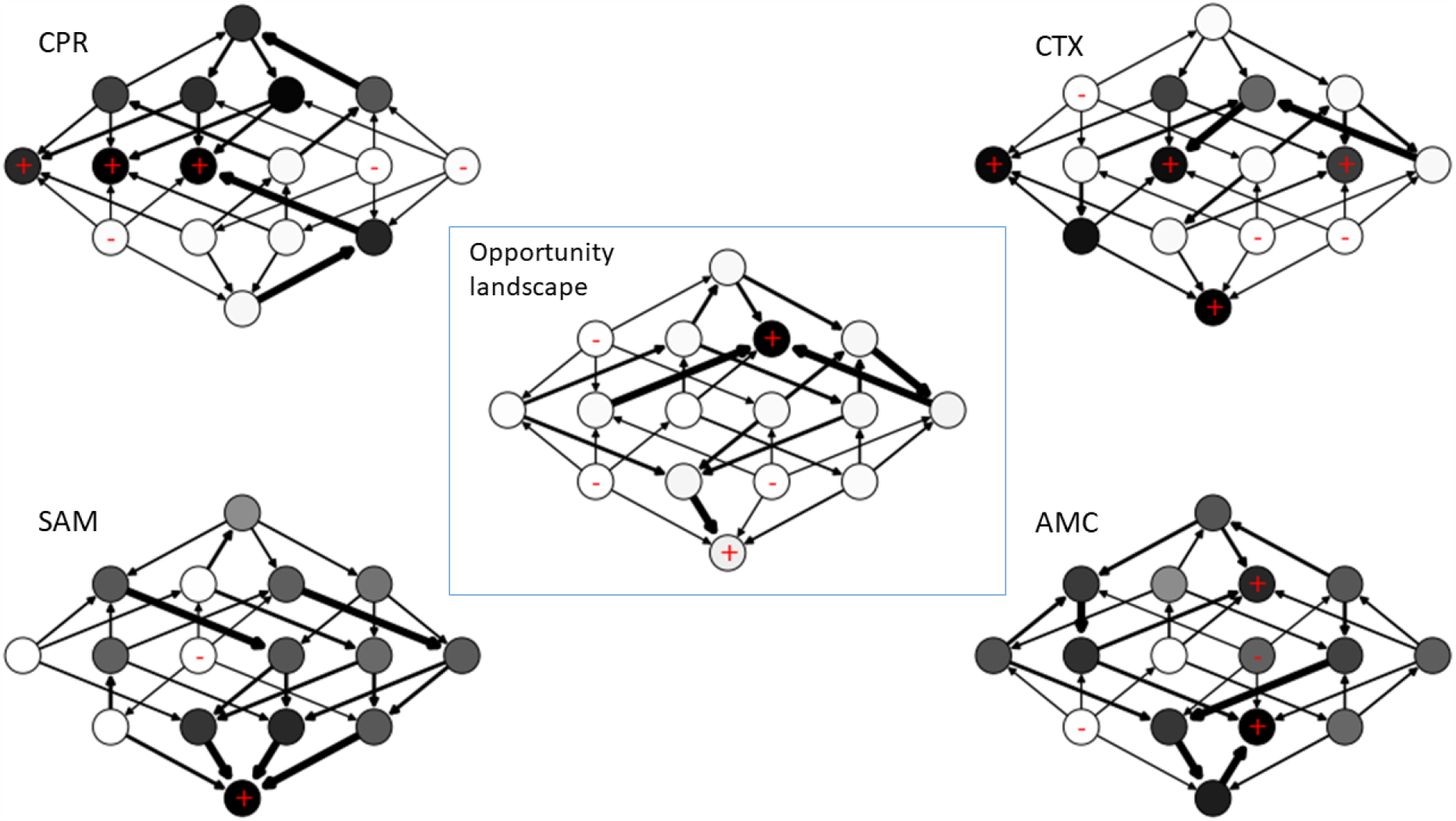
Opportunity landscape for most common policy identified in the RL-genotype condition. As in the MDP-derived policy, just 1/16 genotypes is near a fitness peak in the opportunity landscape.

**Figure S5.**
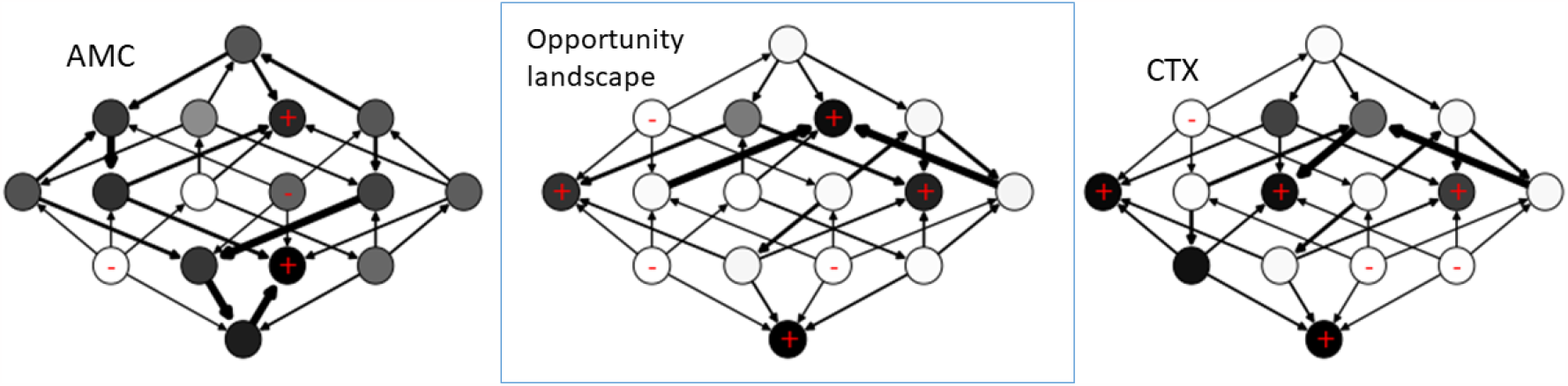
Opportunity landscape for the most common policy identified in the RL-fit condition. The most common RL-fit policy relies on AMC and CTX to control the E. coli population. Assuming the most optimistic combination of these two drug landscapes, 4/16 genotypes are near a fitness peak.

The opportunity landscape for the RL-genotype is almost identical to the opportunity landscape observed for the MDP policy (**Fig S4**). Interestingly, RL-genotype only uses 3 of the 5 drugs in the MDP policy; CPR, CTX, and SAM. RL-fit discovered policies that typically only used two drugs. The most effective RL-fit policies relied heavily on AMC and CTX. We present the resulting opportunity landscape in **Fig S5**. As expected, there are more genotypes with high fitness values under this two-drug paradigm compared to the 4 or 5 drug policies discovered by RL-genotype and the MDP, respectively.

### 1.7 MDP policy

As mentioned in the main text, we computed the MDP policy by formulating a Markov decision process of the strong selection, weak mutation model of evolution under study. We then solved the MDP using backward induction, an algorithm designed to identify an optimal policy for a finite time discrete MDP. The identified policy is a function of current state and current time step, making it even more specific than the policies identified by the reinforcement learning conditions. We show the time and state-specific MDP policy in **Fig S6**. Near the end of an episode (steps 19 and 20), we see a switch to a greedy policy that simply selects the drug with the minimum fitness for a given genotype.

**Figure S6.**
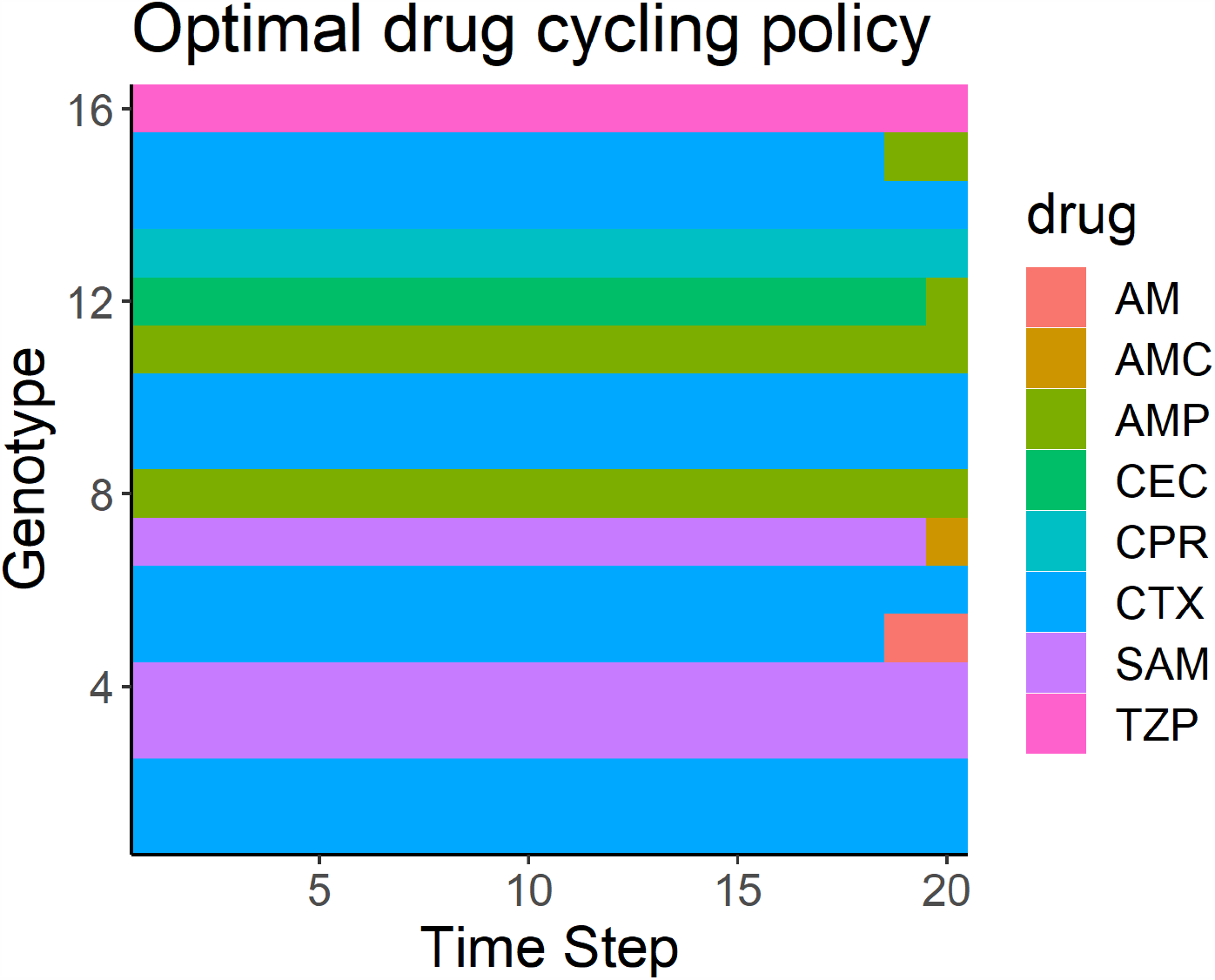
MDP-derived optimal policy for the empirical drug landscapes condition. The X-axis corresponds to numerically encoded genotype while the y-axis corresponds to time step within a given episode. Fill color corresponds to which drug the MDP-derived policy selects for each genotype-time step combination.

We also varied the discount rate (*γ*), between 0 and 1 during the hyperparameter tuning process. In **Fig S7**, we show the effect of gamma on the avergage fitness achieved by the MDP policy. While gamma didn’t have a large effect, likely due to the relatively short length (20 time steps) of each episode, we show that increasing *γ* led to increased performance of the computed MDP policy. We also show that increasing gamma led to increased use of CTX (drug 4).

**Figure S7.**
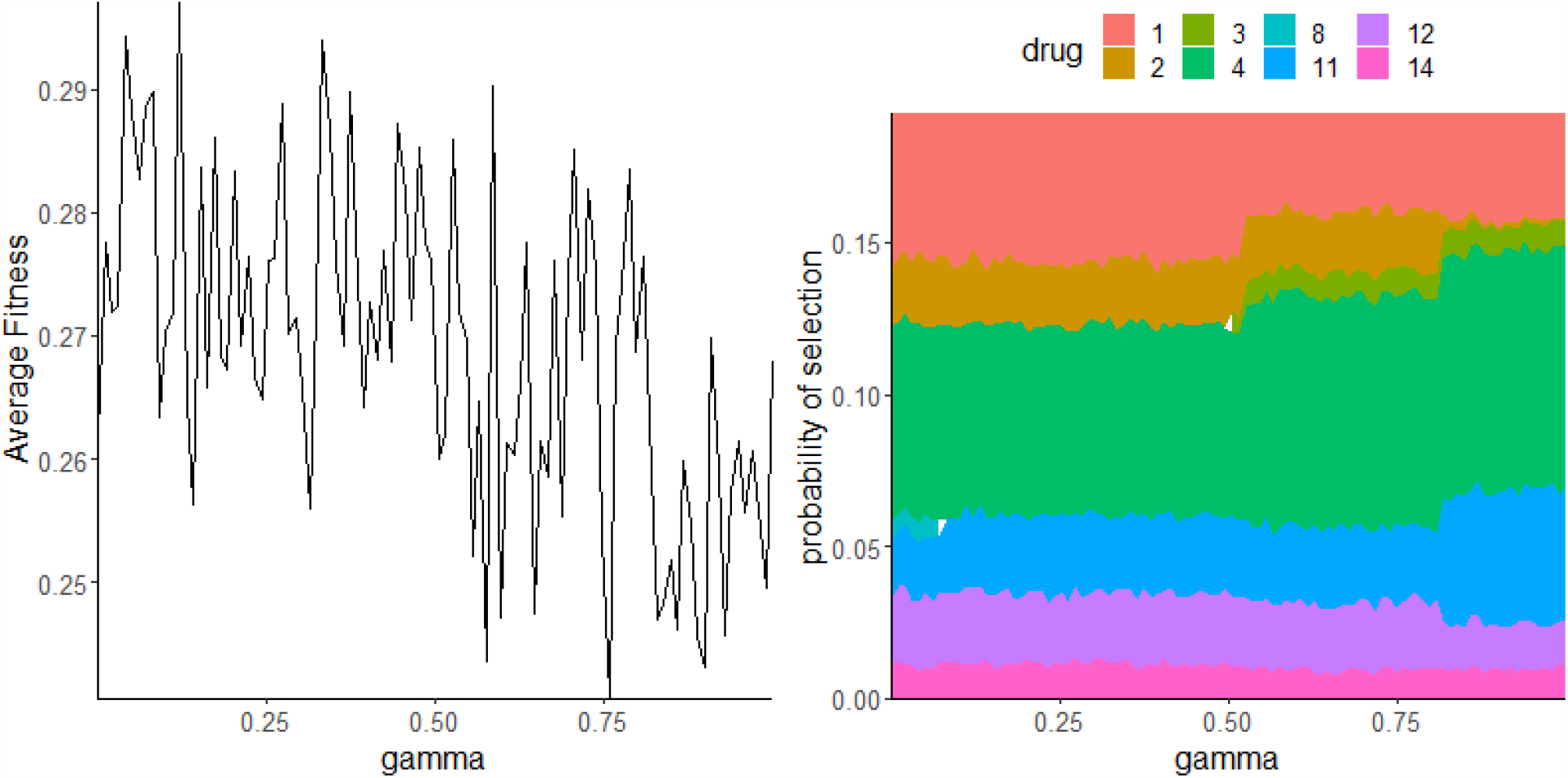
Effect of variation in gamma on optimal policy performance and composition. We show that greedy policies (low *γ*) are slightly less effective compared to non-greedy policies. In panel B, we show how the actual policy changes as *γ* ranges from 0-0.9999.

### 1.8 Additional Analyses

As noted in **Fig 2C** in the main text, we evaluated the performance of all A-B-A-B two-drug cycles to use as a comparison group for RL-fit and RL-genotype. In **Fig S8A**, we examine these combinations in greater depth. We also show the landscape correlation between the two drugs in every combination. We show that anti-correlated landscapes tend to make more effective combinations, likely due to collateral sensitivity. Highly correlated landscapes tend to make ineffective drug combinations, likely due to collateral resistance.

Finally, we evaluated the effect of starting population genotype on the performance of each two-drug combination. We found that the starting genotype of the population had no effect on the overall distribution of performance for these two-drug combinations (**Fig S8**).

**Figure S8.**
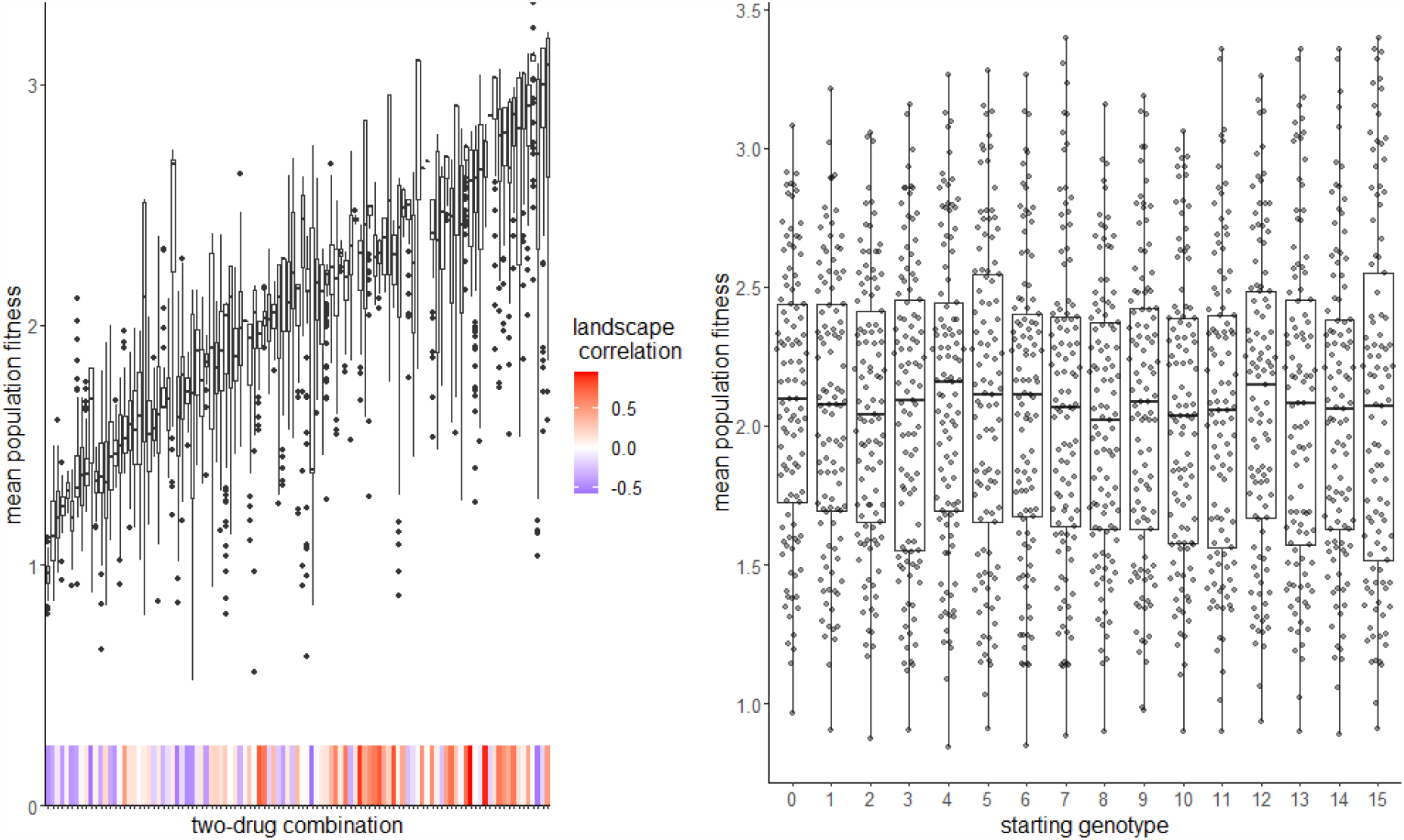
In the left panel, we show the fitness observed under every possible A-B-A-B two drug regime. The heatmap shows the correlation between the two landscapes in a pair, a measure of the expected collateral sensitivity or resistance. In the right panel, we show the effect of starting genotype on the performance of two-drug policies.

